# Microvascular morphology affects radiotherapy outcomes: a computational study

**DOI:** 10.1101/2023.10.16.562646

**Authors:** Luca Possenti, Piermario Vitullo, Alessandro Cicchetti, Paolo Zunino, Tiziana Rancati

## Abstract

Radiotherapy (RT) is the most common cancer treatment, and hypoxia is one of the main causes of resistance to RT. We investigate how microvascular morphology affects radiation therapy results, exploring the role of the microvasculature. Several computational models have been developed to analyze microvascular oxygen delivery. However, few of these models have been applied to study RT and the microenvironment. We generated 27 different networks, covering 9 scenarios defined by the vascular density and the network regularity. Leveraging these networks, we solved a computational mixed-dimensional model for fluid flow, red blood cell distribution, and oxygen delivery in the microenvironment. Then, we simulated a fractionated RT treatment (30 × 2*Gy*_*RBE*_) using the Linear Quadratic model, accounting for oxygen-related (OER) modifications by two different models from the literature. First, the analysis of the hypoxic volume fraction and its distribution reveals a correlation between hypoxia and treatment outcome. The study also shows how vascular density and regularity are essential in determining the success of treatment. Indeed, in our computational dataset, an insufficient vascular density or regularity is sufficient to decrease the success probability for photon-based RT. We also applied our quantitative analysis to hadron therapy and different oxygenation states to assess the consistency of the microvasculature’s role in various treatments and conditions. While proton RT provides a Tumor Control Probability similar to photons, carbon ions mark a clear difference, especially with bad vascular scenarios, i.e., where strong hypoxia is present. These data also suggest a scenario where carbon-based hadron therapy can help overcome hypoxia-mediated resistance to RT. As a final remark, we discuss the importance of these data with reference to clinical data and the possible identification of subvoxel hypoxia, given the size similarity between the computational domain and the imaging voxel.

**Author summary:** The study focuses on investigating the role of microvascular density and morphology in radiation therapy, with particular reference to its role in shaping hypoxia-mediated resistance to treatment. While several computational models have been developed to analyze microvascular oxygen delivery, few have been applied to study RT and the microenvironment. We generated 27 different vascular networks to fill this gap, considering three oxygenation states and three radiation sources (photons, protons, and carbon ions). These cases were analyzed using a complex computational model to describe the state of oxygenation and radiation treatment. The results highlight the importance of vascular density and regularity in determining the success of treatment by shaping hypoxia in the microenvironment. Data suggest that carbon-based hadron therapy can help overcome hypoxia-mediated radiation resistance. As a final remark, these data are also helpful in interpreting possible subvoxel hypoxia with reference to clinical data from different imaging techniques.

## Introduction

Radiotherapy (RT) is the most common treatment for cancer, involving more than half of the patients [1]. Since the first observations in bacteria [2], hypoxia has been known to be a major determinant of tumor resistance to RT, decreasing the damage induced by ionizing radiation [3]. Even if hypoxia is qualitatively defined as a lack of oxygen, a quantitative definition is not unique [4]. Severe hypoxia is often characterized by oxygen concentrations ranging from 0.02% to 0.2% (0.15 to 1.5 *mmHg*), whereas moderate hypoxia referred to oxygen concentrations up to 1% or 2% (7.6 or 15.2 *mmHg*). On the other hand, hypoxia is often classified by its origin as diffusion-limited *chronic hypoxia* or perfusion-limited *acute hypoxia* [3]. The first is related to the distance the oxygen needs to cover to reach the specific tissue area, namely the distance to the closest vessel. The second results from impaired perfusion, reducing the amount of oxygen carried by the microvasculature.

Given the important role of oxygen in RT, some imaging-based methods have been proposed to evaluate hypoxia in tissues, particularly tumors [3, 5, 6]. Such methods take advantage of different technologies such as positron emission tomography (PET), near-infrared spectroscopy (but this is related to vascular oxygen) and magnetic resonance imaging (MRI), which allows the extraction of signals related to the oxygen content of the vascular (BOLD signal) and tissue (TOLD signal) [7]. The identification of hypoxic areas in the tumor, particularly using the TOLD method, is also correlated with the outcome of RT [8]. Further image-based methods have been proposed to identify hypoxic regions by signal combination. For example, Hompand et al.’s method highlights hypoxic regions, estimating the oxygen delivery by the vasculature and the consumption by the tissue [9]. However, we must mention the resolution mismatch between oxygen delivery scale, imaging resolution, and treatment modulation [10]. Indeed, oxygen delivery within the microenvironment occurs at the microscale (hundreds of *µm*), while imaging is characterized by voxels at the *mm* scale [11], and treatment delivery is planned at a slightly larger scale. Such a scale mismatch motivates deeper analyses involving different methods, from computational to *in-vitro*, and *animal* models [12].

The delivery of oxygen to microvessels is usually studied computationally by reducing the number of dimensions of the vascular system through the use of various techniques. From a macroscopic point of view, perfusion has been described using porous media [13–15]. Simultaneously, models that account for the local morphology of the microvasculature have been developed, for example, describing the microvasculature as a collection of concentrated sources, see for example [16] and more recently [17]. This seminal idea has evolved into more advanced multiphysics approaches [18–20], embracing vascular and interstitial flow, red blood cell transport, oxygen transport, and enabling mesoscale analysis of the vascular microenvironment. With such an approach, Powathil et al. validated the prediction of their model against hypoxic markers and *in situ* oxygen measures [21]. New approaches to direct simulation of complex vascular networks with their environment must be performed to be viable for blood flow and oxygen transfer [22]. Such benchmarks can unravel the impact of vascular transport and network morphology on tissue hypoxia in specific applications (e.g., brain [22, 23], tumor [21, 24]).. We recently conducted a thorough sensitivity analysis that demonstrated that the density and structure of a network can have an effect on the oxygen distribution in tissue. [25].

However, few of these models have been applied to study the RT and the microenvironment. Scott and colleagues described the interaction between vascular density and a morphological index [26]. Indeed, they reported variations in correlation between the morphological index and the treatment outcome when considering different vascular densities. In a subsequent study, the same authors showed the importance of microvascular oxygen transport description [27]. They report no influence of vascular morphology on treatment outcomes, but with a constant oxygen content in the vascular network. Schiavo and colleagues generated a wide *cm* vascular geometry by a fractal approach with a hypovascularized core [28]. They studied both chronic and acute hypoxia, showing how model parameters determine hypoxia, varying its position and area. Overall, these studies suggest a key role for the vascular network in determining the outcome of RT.

Two modifying factors must be considered when estimating the RT outcome from computational oxygen distributions. The first one directly regards the oxygen concentration and quantifies the oxygen-mediated radioresistance. The oxygen enhancement ratio (OER) shows the dose scale factor required to obtain the same tumor control when treating hypoxic versus normoxic tumors [29]. The second factor describes the biological action of different ionizing radiations, such as protons and carbon ions [29]. Similarly to the first index, the relative biological efficacy (RBE) is the dose ratio computed referring to the reference radiation (usually photons) and a test one. Hadron therapy utilizes radiations with a Relative Biological Effectiveness (RBE) of 1 or higher, meaning that they have greater biological efficacy than photons. The radiations used in this case are also characterized by a higher linear energy transfer (LET), that is, the energy deposited per unit track, measured by *keV/µm*. Interestingly, the OER is influenced by LET. For example, carbon ions typically have a higher LET than photons, and their effectiveness is less affected by the presence of hypoxia (that is, the OER is lower at a higher LET) [29]. Taking into account hypoxic imaging data, Garrido-Hernandez and colleagues compared different models describing RBE and OER with reference to the treatment plan [30]. They reported some differences in the prediction of the models, but even a significant effect of hypoxia.

Within this framework, our objective was to analyze the effect of microvasculature on RT treatment using a state-of-the-art computational model. On the basis of these results, we address the influence of microvascular morphology (vascular density and regularity) on RT outcome, to understand the role of the microvasculature. Furthermore, we show how this approach allows for quantitative analysis of the microvasculature and the microenvironment with reference to RT. Lastly, we extend such analysis to hadron therapy and altered oxygenation states, so that we assess whether the role of microvasculature is consistent through different types of treatment and under different conditions.

## Materials and methods

The workflow of this study comprises three steps: (i) generating different configurations of the geometry of the microvascular network, (ii) computing the oxygen concentration in the microvascular environment, and (iii) applying radiobiological models to estimate the result of radiotherapy. In the following paragraphs, we detail the methods for each of these steps. The results are then analyzed and visualized by Paraview [31] and Prism [32].

### Generation of microvascular networks

We consider a region of 1 *mm* × 1 *mm* × 150*µm* that describes a ideal portion of vascularized tissue. To generate representative vaacular networks in that region, we adopt the strategy we previously proposed [25]. This method uses Voronoi tessellation on a plane, resulting in a space-filling network following biomimetic principles [18, 33] and comprising bifurcation and anastomosis alone. As Voronoi networks are uniquely defined by a set of seed points, to initialize the network generation, we adopt a two-step approach based on the placement of seed points in a square 1 *mm* × 1 *mm*, subdivided into *positive* and *negative* points, with opposite roles. First, we uniformly distribute a set of positive seeds in the plane. Then, a set of negative seeds is generated based on a Poisson distribution centered in (0.5 *mm*, 0.5 *mm*). Each negative seed deletes a positive one, leading to a heterogeneous vessel distribution. Finally, a positive seed is added in the center of the square. Thanks to this method, we have produced networks characterized by different metrics. In particular, we control the surface-to-volume ratio *S/V* (that is, the lateral surface of the network over the volume of the tissue) by controlling the positive seeds. On the contrary, the irregular morphology, quantified by the maximum distance of any point in the network (*d*_*max*_), is determined by the ratio among the negative and positive seeds. Finally, we generate a quasi-3D network by moving the junctions and the boundary nodes along the direction perpendicular to the plane, to fill the thickness of 150 *µm*.

We generate a set of 27 networks, representing 9 different conditions in triplicates (Fig. S1). The nine conditions combine three values of *S/V* and three values of *d*_*max*_, ranging from poor to highly vascularized tissues and from regular to irregular morphology (Table 1).

**Table 1.** Values for the parameters of the model. The values of the parameters are here reported, along with their units and references. For the oxygen scenarios, the parameters specified were changed, while the others are kept constant.

| Network morphology |  |  |  |
| --- | --- | --- | --- |
| Symbol | Parameter | Value | Ref. |
| $S/V$ | Vascular area over tissue volume | $3500 - 7000 - 14000 \text{ m}^{-1}$ | [34] |
| $d_{max}$ | Maximum distance from the network | $120 - 160 - 240 \text{ }\mu\text{m}$ | - |
| Fluid problem $\mathcal{F}$ | | | |
| $K$ | Tissue permeability | $3.6 \times 10^{-17} \text{ m}^2$ | [35] |
| $\mu_t$ | Interstitial fluid viscosity | $1.2 \times 10^{-3} \text{ Pa s}$ | [36] |
| $\mu_v$ | Blood viscosity | computed | [37] |
| $R$ | Vascular radius | variable | [25] |
| $Lp$ | Vascular wall hydraulic conductivity | $1.4 \times 10^{-10} \text{ m}^2 \text{ s/kg}$ | [38] |
| $\pi_v - \pi_t$ | Transmural osmotic difference | $640 \text{ Pa}$ | [17, 39] |
| $\sigma$ | Reflection coeff. for proteins | 0.82 | [35] |
| Oxygen problem $\mathcal{O}$ | | | |
| $HF$ | Hüfner factor | $1.36 \text{ ml}_{O_2}/g_{Hb}$ | [40] |
| $MCHC$ | Mean Corpuscular Hematocrit Concentration | $0.34 \text{ g}_{Hb}/\text{ml}_{RBC}$ | [40] |
| $\eta_{pl}$ | Oxygen solubility in plasma | $2.82 \times 10^{-5} (\text{ml}_{O_2}/\text{ml})/\text{mmHg}$ | [41] |
| $p_{s50}$ | $pO_2$ at hemoglobin half-saturation | $27 \text{ mmHg}$ | [40] |
| $\gamma_{Hb}$ | Hill exponent | 2.64 | [40] |
| $V_{max}$ | Maximum oxygen consumption rate | $2.47 \times 10^{-4} (\text{ml}_{O_2}/\text{s})/\text{cm}^3$ | [40] |
| $\eta_t$ | Oxygen solubility in the tissue | $3.89 \times 10^{-5} (\text{ml}_{O_2}/\text{cm}^3)/\text{mmHg}$ | [41] |
| $p_{m50}$ | $pO_2$ at half consumption rate | $0.5 \text{ mmHg}$ | [40, 41] |
| $P_{O_2}$ | Microvascular wall permeability | $3.5 \times 10^{-5} \text{ m/s}$ | [42] |
| $\varepsilon_t$ | Oxygen diffusion coefficient in the tissue | $2.41 \times 10^{-9} \text{ m}^2/\text{s}$ | [41] |
| $\varepsilon_v$ | Oxygen diffusion coefficient in the vasculature | $2.18 \times 10^{-9} \text{ m}^2/\text{s}$ | [41] |
| Boundary conditions |  |  |  |
| $p_v\vert _{in}$ | Pressure at vascular inlets | $40 \text{ mmHg}$ | [43] |
| $p_v\vert _{out}$ | Pressure at vascular outlets | $15 \text{ mmHg}$ | [43] |
| $H_{in}$ | Discharge hematocrit at vascular inlets | 0.45 | [40] |
| $pO_2\vert _{in}$ | $pO_2$ at vascular inlets | $90 \text{ mmHg}$ | [44–46] |
| Radiotherapy |  |  |  |
| $N$ | Tumor cells in the reference volume | $15k$ | [47] |
| $a_1$ | WEN parameter | $0.22 \text{ Gy}^{-1}$ | [48] |
| $a_2$ | WEN parameter | $0.0024 \text{ Gy}^{-1}(\text{keV}/\mu\text{m})^{-1}$ | [48] |
| $a_3$ | WEN parameter | $0.05 \text{ Gy}^{-1}$ | [48] |
| $a_4$ | WEN parameter | $0.0031 \text{ Gy}^{-1}(\text{keV}/\mu\text{m})^{-1}$ | [48] |
| $b_1$ | WEN parameter | $0.4 \text{ Gy}^{-1}$ | [48] |
| $b_2$ | WEN parameter | $0.015 \text{ Gy}^{-1}$ | [48] |
| $k_{RT}$ | $pO_2$ for half radiosensitivity (WEN) | $3 \text{ mmHg}$ | [48] |
| $\gamma$ | TIN parameter | 1.38 | see Appendix |
| $M$ | TIN parameter | 2.81 | see Appendix |
| $a$ | TIN parameter | $522.45 \text{ keV}/\mu\text{m}$ | see Appendix |
| $b$ | TIN parameter | $1.24 \text{ mmHg}$ | see Appendix |
| Oxygen scenarios |  |  |  |
| Symbol | Parameter | Value | Scenario |
| $p_v\vert _{in}$ | Pressure at vascular inlets | $30 \text{ mmHg}$ | Acute hypoxia |
| $p_v\vert _{out}$ | Pressure at vascular outlets | $25 \text{ mmHg}$ | Acute hypoxia |
| $V_{max}$ | Maximum oxygen consumption rate | $4.94 \times 10^{-4} (\text{ml}_{O_2}/\text{s})/\text{cm}^3$ | High consumption |

### Modeling blood flow and oxygen transfer

The microvascular environment model used in this work belongs to the mesoscale approach family [49]. We describe the microenvironment as a three-dimensional domain, Ω, in which we embed the microvascular network represented as a collection of one-dimensional channels, i.e., the vascular domain is a metric graph denoted by Λ. In particular, the coupling of models at different dimensions requires specific mathematical methods [50, 51]. However, it brings some significant advantages at the computational level that will be discussed later on.

In these two domains, we assembled three mathematical problems that describe different phenomena: (i) fluid flow F [50], (ii) transport of red blood cells H [18], (iii) transport of oxygen 𝒪 [20].

### Microvascular flow and red blood cells transport

First, we model fluid flow within the vasculature and surrounding tissue (ℱ). Briefly, an interstitial fluid is described as a Newtonian fluid in a porous medium, employing Darcy’s equation. On the other hand, we use the Poiseuille equation for blood flow in the vasculature. The effects of red blood cells (ℋ) are included in the model considering their effect on viscosity, by the Fahraeus-Lindqvist effect [37], and their splitting when traveling in a bifurcation, namely the Zweifach-Fung effect [52]. Fluid filtration through the microvascular wall is described by the classical Starling equation, which accounts for hydraulic and osmotic pressures. The problem is complemented by the mass conservation equations in the two domains (the tissue Ω and the vasculature Λ), and it reads as follows:

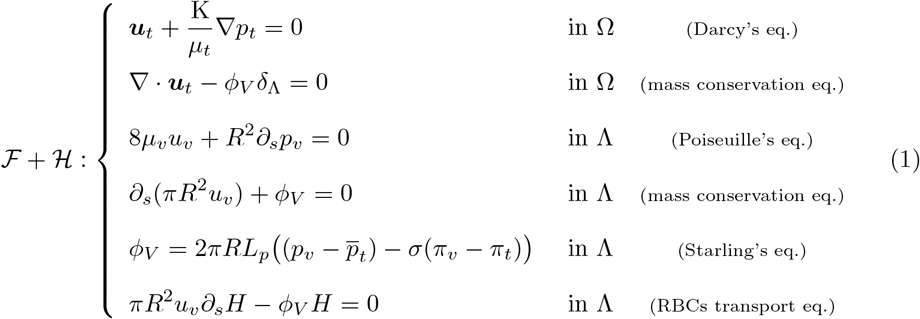

where the subscripts *t* and *v* represent the tissue and vasculature, respectively, *u* is the fluid velocity, *p* is the fluid pressure, *µ* is the viscosity, K is the porous medium permeability, *R* is the radius of the vessel, *ϕ*_*V*_ fluid flow across the vascular wall affected by the vascular hydraulic conductivity, *L*_*p*_, the reflection coefficient for proteins, *σ* and the osmotic pressure, *π*. Further details regarding modeling equations, procedures to handle network junctions, and numerical solutions of the problem can be found in [18, 53]. The problem is then defined by considering boundary conditions, setting variable values for the inlet and outlet pressures, and hematocrit inlet. For tissue, homogeneous conditions are enforced so that no flow can leave the domain.

### Oxygen transport

The oxygen transport equations (𝒪) for the two domains comprise diffusion, consumption, and advection, using data from the models ℱ and ℋ in a one-way coupling scenario. In addition to the above-mentioned component, we introduce three particular phenomena: (i) oxygen binding to hemoglobin, (ii) oxygen uptake dynamics, and (iii) oxygen exchanges through microvascular walls.

For the first, we recall that oxygen is present in the system as solved in plasma/interstitial fluid (*C*_*v*_ or *C*_*t*_) or bound to hemoglobin 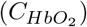. The total oxygen content is the sum of the two in the vasculature, that is, the region containing RBC and hemoglobin. We assume the binding dynamics as an instantaneous process, described by the Hill’s equation:

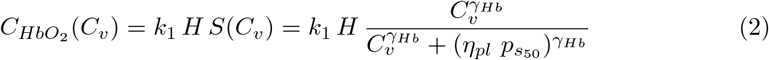

where *k*_1_ is a constant value determined by the product of the Hüfner factor (*HF*) describing the oxygen binding capacity of human hemoglobin and the Mean Corpuscular Hematocrit Concentration (*MCHC*). The remaining parameters of (2) are the oxygen solubility in plasma *η*_*pl*_, the oxygen partial pressure at half-saturation of hemoglobin 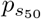, and the Hill exponent *γ*_*Hb*_.

Secondly, we describe the cell uptake in the tissue by the well-known Michaelis-Menten equation [54]:

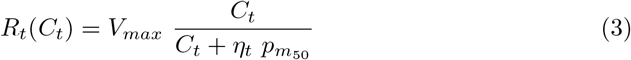

where *V*_*max*_ is the maximum oxygen consumption rate, *η*_*t*_ is the oxygen solubility in the tissue, and 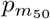 is the Michaelis-Menten constant, i.e. the oxygen partial pressure at half consumption rate.

The oxygen exchanges include diffusion and advection through the microvascular wall, as a semi-permeable membrane [55]:

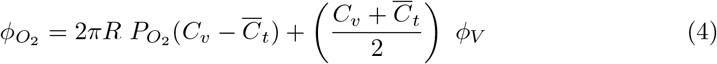

where 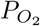 is the microvascular wall permeability.

Comprehensively, the oxygen transport model reads as follows:

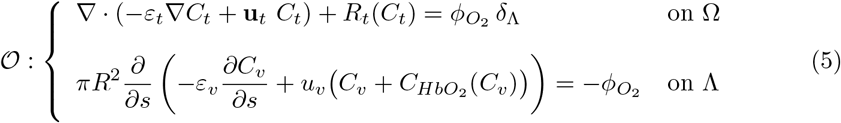

where *ε*_∗_, ∗ = *v, t* are the diffusion coefficients in the blood and tissue compartments. The model describes oxygen transport through the microvasculature and delivery to the tissue through flux 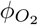. We note that this is a mixed-dimensional mathematical formulation that combines a 3D region, Ω with a 1D network Λ. At the mathematical level, the coupling between the two is provided by a Dirac measure distributed along the network, here denoted by *δ*_Λ_. Eventually, oxygen reaches the cells and cell uptake occurs. The problem is complemented by boundary conditions that define oxygen concentrations at network inlets, the null derivative at network outlets, and null concentration gradients at the tissue faces, i.e., no diffusive flux. The complete details related to model derivation, discretization, solving, and validation are reported in [20].

### Simulations of oxygen transport in the microenvironment

For complex geometrical configurations, explicit solutions to problems ℱ, *ℋ, 𝒪* are not available. Numerical simulations are the only way to apply the model to real cases. The discretization of these models, described in [18], is achieved by the finite element method (Appendix. S1). Because of the particular mathematical structure of the model, based on mixed-dimensional differential equations, there is no commercially available simulator that can handle it. All simulations have been performed using an internal C++ code based on the GetFem++ open source library [56].

The main advantage of the mixed-dimensional formulation adopted here is that the discretizations of the equations defined in the tissue and in the vascular network are completely independent in terms of the computational grids and the numerical schemes. We discretize the branches of the vascular network as separate subdomains. Each of them is approximated by a piecewise straight line. The problems of blood flow and oxygen transfer are approximated using continuous piecewise polynomial finite elements. The interstitial flow problem is approximated using mixed finite elements.

Because of the non-linear relationships present in the constitutive laws of the model, the problem has been linearized, and the solution of the coupled problem has been reached via an iterative process. At each iteration, the numerical discretization schemes described above provide a high-dimensional linear system of equations, solved by means of state-of-the-art iterative (linear) solvers with suitable preconditioners. For further details on the computational methods, we refer to [18, 42, 50, 51].

The ideal tissue slab with dimension of 1 *mm* × 1 *mm* × 150*µm* has been discretized by means of a uniform tetrahedral mesh of 40 × 40 × 6 nodes on each side, consisting of 57600 tetrahedra. This numerical resolution has been considered satisfactory after performing a mesh sensitivity analysis. In particular, the mean relative error in the *L*^2^ norm on the approximation of the oxygen partial pressure between the considered mesh and a new one with higher resolution (precisely 50 × 50 *times* 8 points per side) resulted in less than 3%.

The simulation of each scenario of blood flow and oxygen transfer discussed below is not computationally demanding on a multi-core processor with good performance. However, scaling up these computations to the whole tumor scale will certainly require high-performance computing platforms.

### Radiobiological models for the radiotherapy outcome

We model radiotherapy treatment using the linear-quadratic (LQ) model, which is the most used radiological model. It is based on two different parameters that describe the radiosensitivity of cells or tissue. The first parameter *α* describes the lethal damage resulting from a ‘single’ hit, whereas *β* is related to ‘multiple’ hits, namely the interaction of multiple radiation tracks [57]. Combining these two parameters and the dose delivered (*D*), the surviving fraction (*SF*) is estimated as follows:

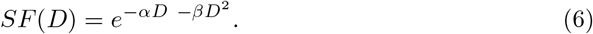

The admissible values of *SF* range from 0 to 1, representing the fraction of cells that survived treatment with the specified dose *D*. In current clinical practice, the total dose of treatment is divided into fractions *n*_*f*_, usually administered daily, that is, fractionated treatments [1]. To describe such a treatment schedule, the cumulative effect of each fraction is described by a slightly modified LQ model, by applying the LQ model recursively:

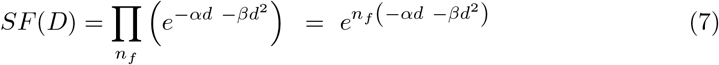

where *d* = *D/n*_*f*_ is the fraction dose. We remark that tumor growth during treatment is not included in this equation.

Due to the small tissue region analyzed, we assume a constant dose *D* over the volume. The same assumption is not valid for the oxygen content, which may differ throughout the region, affecting treatment results. To account for the oxygen effect, we adopt two different models: the Wenzl model (WEN) [48, 58] and the Tinganelli model (TIN) [59]. These are basically modifications of the classical LQ model. The model proposed by Wenzl and colleagues defines the variation of *α* and *β* with the oxygen partial pressure and the linear energy transfer (LET) of ionizing radiation:

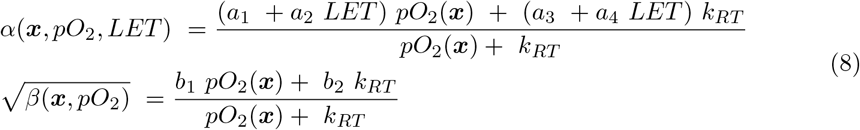

where *k*_*RT*_ is the oxygen tension for radiosensitivity equal to half of its maximum value. Expressions of *α*(***x***, *pO*_2_, *LET*) and *β*(***x***, *pO*_2_) are inserted directly into the LQ equation:

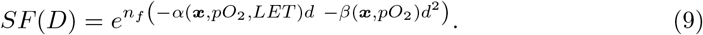

On the contrary, the Tinganelli model is based on the definition of the Oxygen Enhancement Ratio (OER), which is then used to scale the dose:

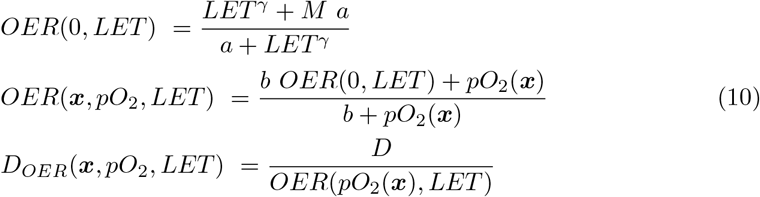

where *M, a, γ, b* are parameters fitted to experimental data, and *D*_*OER*_(***x***, *pO*_2_, *LET*) is the dose corrected for the oxygen effect, to be included in the LQ model. Therefore, TIN’s *α* and *β* do not change with LET or *pO*_2_, and the resulting model reads as follows:

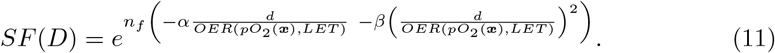

To properly compare the models, we used the parameters of the WEN model published in [48], and fit the TIN parameters to the data reported by WEN and colleagues. Further details on the parameter and the fitting procedure are described in Appendix. S1.

These two LQ-based models allow us to compute the *SF* (***x***). Based on that, we calculate the tumor control probability (*TCP*), which describes the probability of successful treatment. It is based on the number of cells surviving the treatment (SF) and the initial number of clonogenic cells *N* :

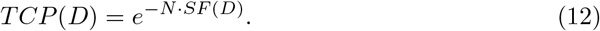

In particular, since we analyze the spatial variability of the RT response on the microscale, we describe *SF* as a spatially dependent map *SF*(*D*, ***x***). We employ the methods presented in [60] to compute the *TCP* (*D*) from the *SF* distribution, leveraging the Poisson process assumption and the ‘parallel-like’ behavior of the tumor (i.e. even a small portion of the tumor surviving results in tumor relapse, and therefore TCP is low):

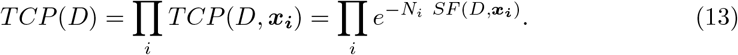

In summary, we obtain *pO*_2_(***x***) in the tissue solving the ℱ, ℋ, and 𝒪. From this we compute *SF* (***x***) by applying the WEN and TIN models. Finally, we consider the spatial distribution *SF* (***x***) to obtain a probability value for *TCP* (*D*) (equation 13).

### Simulated treatment schedule and sources

We initially define the treatment schedule for photons, which is the most common radiation therapy treatment. The total dose of treatment is 60 *Gy*, divided into 30 fractions at 2 *Gy* per fraction, that is, a treatment 30 × 2*Gy*. We point out that this treatment is compatible with the clinical schedules and the clinical dose received by the tumors. For example, non-small cell lung cancers receive 30 × 2*Gy* [61], and prostate cancer patients are usually treated with 60 − 72 *Gy* in fractions 20 − 30 [62]. In our setting, 30 × 2*Gy* treatment guarantees a highly successful treatment (*TCP* ≃ 1) in well-oxygenated tumors (*pO*_2_ *>* 5 *mmHg*).

Furthermore, we analyze a similar treatment schedule for hadron therapy, particularly in the case of protons and carbon ions. These different sources have different biological effects. To consider this, the dose is usually scaled by *RBE* [29]. As a consequence, two different doses are defined: the physical dose, that is, the real dose physically measured, and the biological dose scaled by *RBE*. The biological dose is then used in the LQ models. We compare different sources considering the same biological dose (2 *Gy*_*RBE*_ per fraction), namely, the physical dose is 2*/RBE Gy* per fraction. Furthermore, photons and hadrons also differ in *LET* (photons: 2 *keV/µm*; protons: 4 *keV/µm*; carbon ions: 75 *keV/µm* [29, 63]), which enters in radiobiological models and determines the response to hypoxia.

#### Oxygenation scenarios

We calculate the oxygenation of the microenvironment, namely *pO*_2_ in the domain Ω, under three different conditions. The first is the reference conditions with parameters describing a tumor tissue with no particular characteristics (see Table 1). On the contrary, the second and the third scenarios are alterations of such a reference condition. In particular, the second scenario describes a lower oxygen delivery due to reduced blood flow. Such perfusion-limited hypoxia has been known for years as acute hypoxia (AH) [3, 64]. On the other hand, the third scenario describes an increased oxygen demand by the tumor, so that the oxygen consumption is greater. Such a condition intensifies the possible chronic hypoxia present in the reference scenario. Indeed, such hypoxia is related to diffusion and, therefore, related to the distance of a specific area from the closest vessel [3, 64]. Increased oxygen consumption reduces the distance that oxygen can reach by diffusion before uptake by cells occurs.

## Results

We compute the RT outcome referring to different sources (photons, protons, and carbon ions) and different states of oxygenation in the microenvironment. The results are presented starting from the classical photon-based RT. Then, hadron therapy is analyzed, also considering non-standard oxygenation conditions.

### Oxygen distribution varying microvascular density and morphology

Using the numerical simulations of the models ℱ, ℋ, and 𝒪, we obtain the oxygen distributions in the tissue domain Ω (Fig. 1). The resulting oxygen content agrees with the ranges described in the literature, which anyway differs between tissues, due to differences in vascularization and tissue oxygen demand [5, 65]. When considering a regular network, tissue oxygenation increases with microvascular density, as is evident from the first row in panel 1a, and is quantified by graph 1b (blue columns referring to the lower *d*_*max*_). The same trend is also reported for irregular networks, namely, for higher *d*_*max*_ values. In contrast, the increase of *d*_*max*_ effectively reduces the average partial oxygen pressure in medium and highly vascularized cases only. In the low-vascularized case, the regularity of the network has a little to no effect.

**Figure 1.**
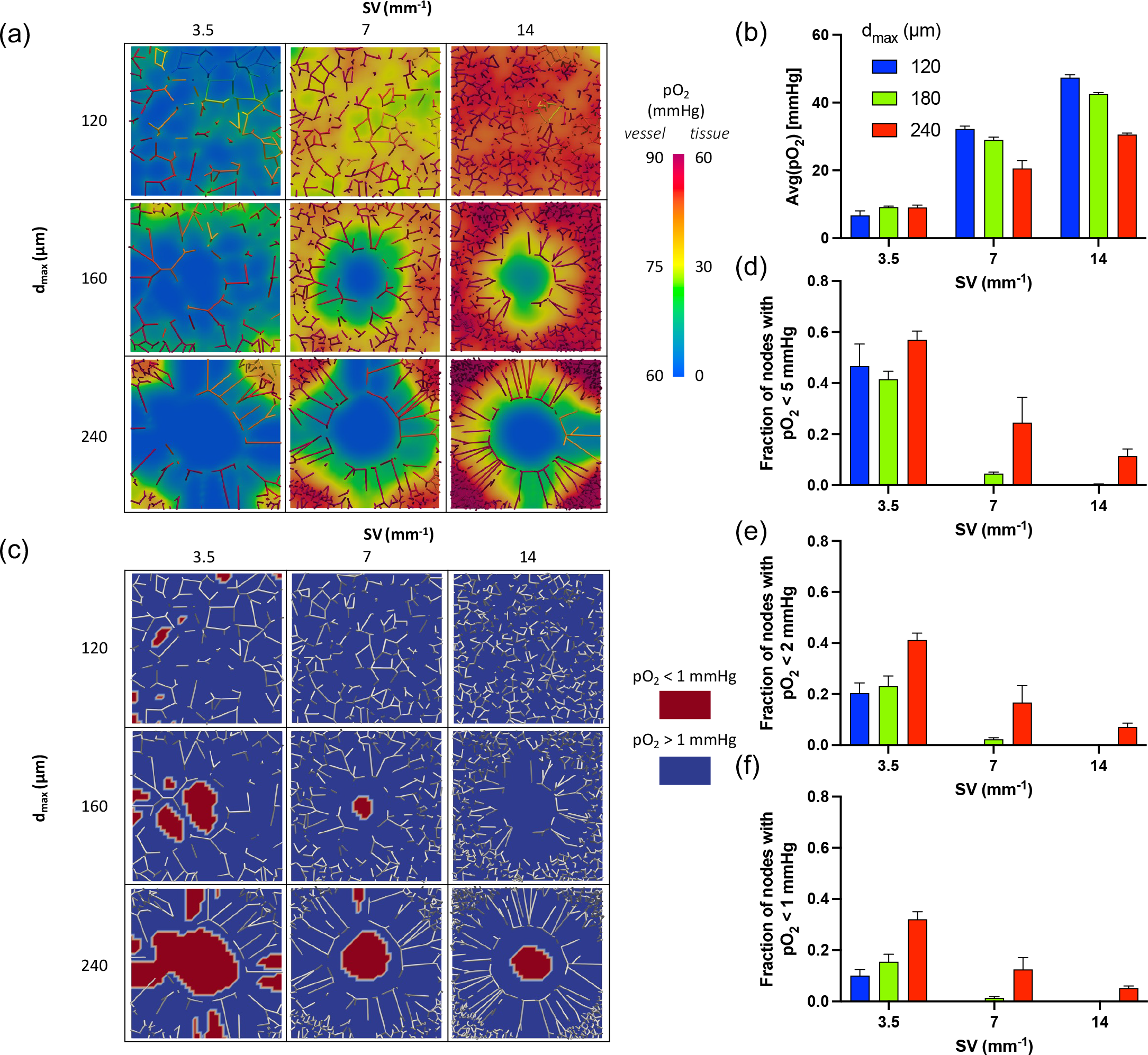
Oxygen distribution in the nine cases for the reference scenario. (a) Oxygen partial pressure in the tissue Ω and microvascular Λ domains. A single replicate of the nine cases (3 *S/V* × 3 *d*_*max*_) is shown. (b) Average oxygen partial pressure in the tissue in the nine cases considered. (c) Localization of low oxygen areas, setting a threshold (*th*) equal to 1 *mmHg*. Similar maps for *th* = 2*mmHg* and *th* = 5*mmHg* are available in the Fig. S2. The other panels show the fraction of nodes below the threshold in the nine cases, setting the threshold to 5 *mmHg* (d), 2 *mmHg* (e), and 1 *mmHg* (f).

On the other hand, considering the low oxygen volume fractions (i.e., node fraction), the irregularity of the network plays a role in all the cases analyzed (Figs. 1 cf. and Fig. S2). In general, the more irregular the network is, the larger the low oxygen fraction is. Additionally, a denser microvascularization results in smaller low-oxygen fractions.

We remark the presence of hypoxic volume in the case *S/V* = 14*mm*^−1^, *d*_*max*_ = 240*µm*. Indeed, our model identifies regions with low oxygen content in highly vascularized tissues if the network morphology is not regular enough. Generally, a higher grade of irregularity (*d*_*max*_) is required to produce hypoxia in higher vascularized tissues. For example, the hypoxic volume appears in the case *S/V* = 7*mm*^−1^, *d*_*max*_ = 160*µm*, but not in *S/V* = 14*mm*^−1^, *d*_*max*_ = 160*µm*.

### Computing TCP with photons RT

Based on the results shown in the previous section, we computed the surviving fraction *SF* (***x***) using the two radiobiological models and the corresponding tumor control probability (Fig. 2). Starting with the WEN model, three cases have TCP →1, namely, the treatment is successful in these cases (*S/V* = 7*mm*^−1^ - *d*_*max*_ = 120*µm*, and *S/V* = 14*mm*^−1^ - *d*_*max*_ = {120; 160} *µm*). These cases are characterized by a tiny or null hypoxic fraction (Fig. 1 c-f), which combines good vascularization with regular microvascular morphology. In fact, even a small hypoxic volume (considering the strong definition with *pO*_2_ ≤ 1 *mmHg*) produces an evident drop in TCP, as can be seen in the cases *S/V* = 7*mm*^−1^ - *d*_*max*_ = 160*µm*. Furthermore, the highly irregular morphology of the network reduces the TCP in all conditions, as shown by the TCP drop when considering *S/V* = 14 *mm*^−1^ cases.

**Figure 2.**
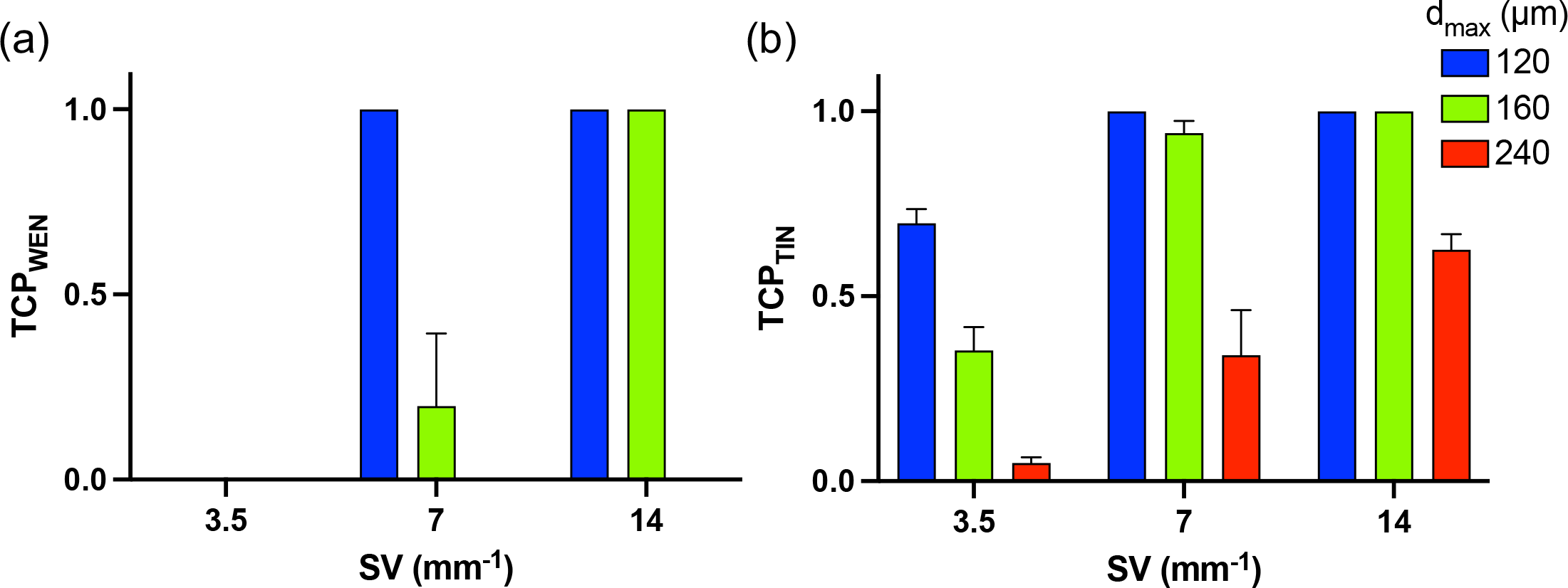
Computed TCPs for the nine cases with photons RT. Resulting TCPs referred to 30 × 2*Gy* treatment with photons RT using the WEN (a) and the TIN (b) models applied to the 9 cases (3 *S/V* × 3 *d*_*max*_). TCP → 1 depicts successful treatments.

The TIN model is characterized by a less sharp response to hypoxia. Consequently, the TCP estimates are higher than the WEN ones. However, the interpretation of the results is similar, with few successful treatments (TCP →1). The main difference is observed in the case *S/V* = 7*mm*^−1^, *d*_*max*_ = 160*µm*, which has a TCP closer to 1 with the TIN model. Such a smoother response to hypoxia shows two additional points. First, the data show that greater vascularization generally improves TCP, while morphological irregularities have a detrimental effect on the outcome of the treatment. Second, the TCP for the cases *S/V* = 14*mm*^−1^ - *d*_*max*_ = 240*µm* and *S/V* = 3.5*mm*^−1^ - *d*_*max*_ = 120*µm* are similar. Therefore, bad microvascular morphology can determine a bad outcome of treatment, even in highly vascularized tissues, and a similar result can be achieved with a regular low-density microvascular network.

### Comparing TCP from photons and hadron therapy

We analyze the same nine cases treated with protons and carbon ions using the same biological dose and schedule (Fig. 3). First, proton-based RT has outcomes similar to those of photon treatment. Therefore, considerations of microvascular architecture and morphology still hold for proton therapy, determining the outcome of the treatment. Again, increased vascularization and morphological regularity improve TCP. We remark that we are using the same biological dose and, consequently, a similar outcome is expected. However, they differ slightly. This is due to the different *LET* associated with the two sources: 2 *keV/µm* for photons and 4 *keV/µm* for protons [29]. *LET* determines also the response to hypoxia, as can be seen in equations 8 and 10. In particular, ionizing radiation characterized by greater *LET* results in being less sensitive to hypoxia-mediated radioresistance. Therefore, they result in a greater TCP. This effect is slightly noticeable comparing photons and protons but is evident when considering carbon ions (*LET* = 75 *keV/µm*) [29, 63]. As a result, carbon ions are effective in treating all of the cases analyzed, without influence on microvascular density and morphology.

**Figure 3.**
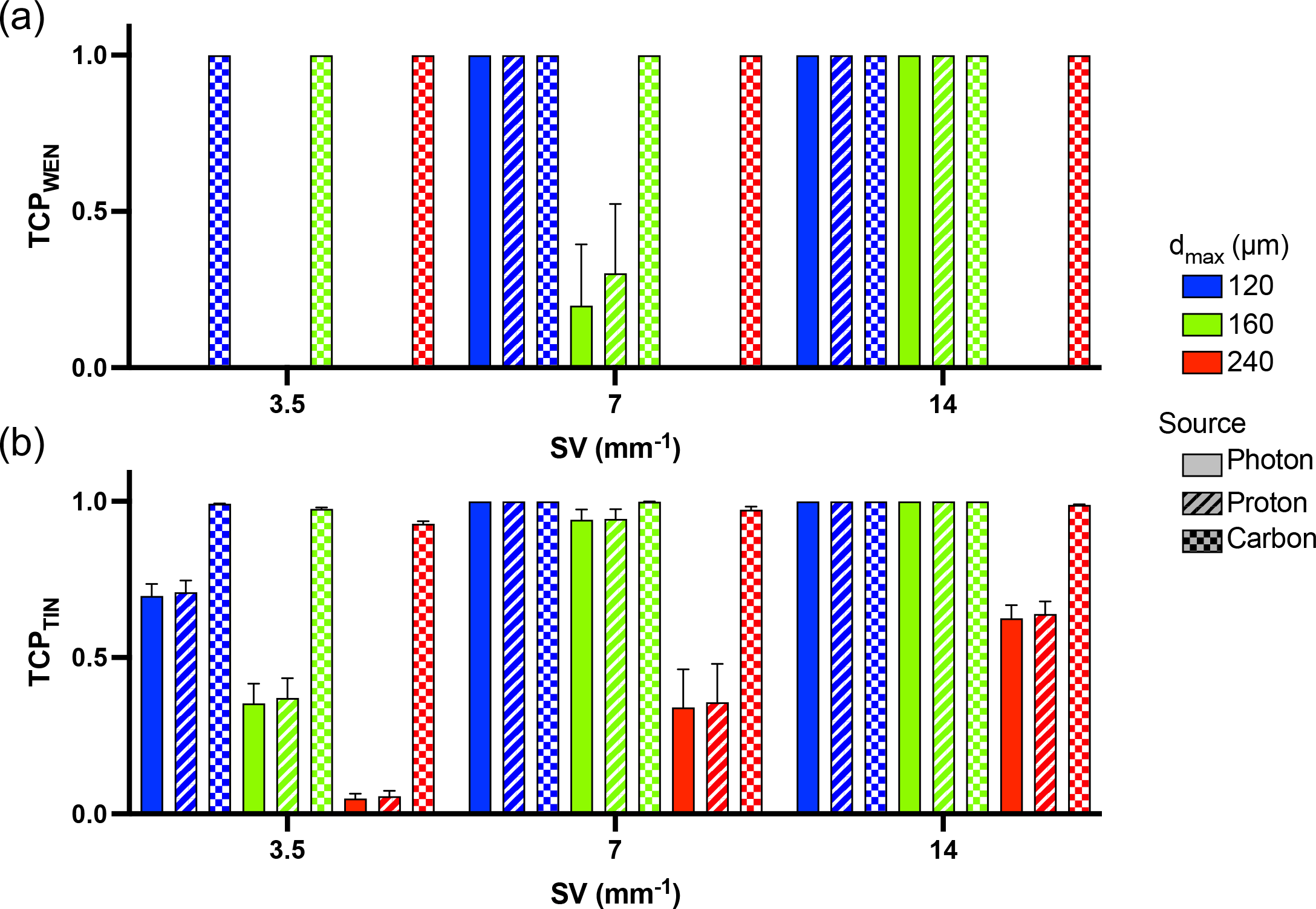
Computed TCPs for the nine cases with three different sources: photons, protons, and carbon ions. Resulting TCPs referred to 30 × 2*Gy*_*RBE*_ treatment using the WEN (a) and the TIN (b) models. RT treatment is simulated considering photons, protons, and carbon ions applied to the 9 cases (3 *S/V* × 3 *d*_*max*_). TCP → 1 depicts successful treatments.

### Influence of acute hypoxia and tumor oxygen consumption

We tested the same nine vascular conditions that reduced the vascular flow rate and increased the tumor oxygen demand, keeping all other parameters at their reference values. First, we address oxygenation in the tissue (Fig. 4). Both altered scenarios decrease the average oxygen content in the tissue with respect to the reference state. Among the two, increased consumption induces a stronger reduction, according to previous work on oxygenation [20, 66, 67]. An important difference between the two scenarios is how they act in the two domains. Accute hypoxia first acts on the microvascular network, leading to a more pronounced decrease of *pO*_2_ in the network (Fig. 4a compared to the left columns). On the other hand, an increase in consumption reduces the oxygen concentration of the tissue uniformly, with lower effects on the network. Such a higher oxygen consumption increases the microvascular density required to guarantee good oxygenation. In fact, the levels in the average partial oxygen pressure for cases *d*_*max*_ = {120, 160, 240} *µm* - *S/V* = 7*mm*^−1^ - *High consumption* are similar to those of *d*_*max*_ = {120, 160, 240} *µm* - *S/V* = 3.5*mm*^−1^ - *Reference*, depicting an insufficient microvascular density.

**Figure 4.**
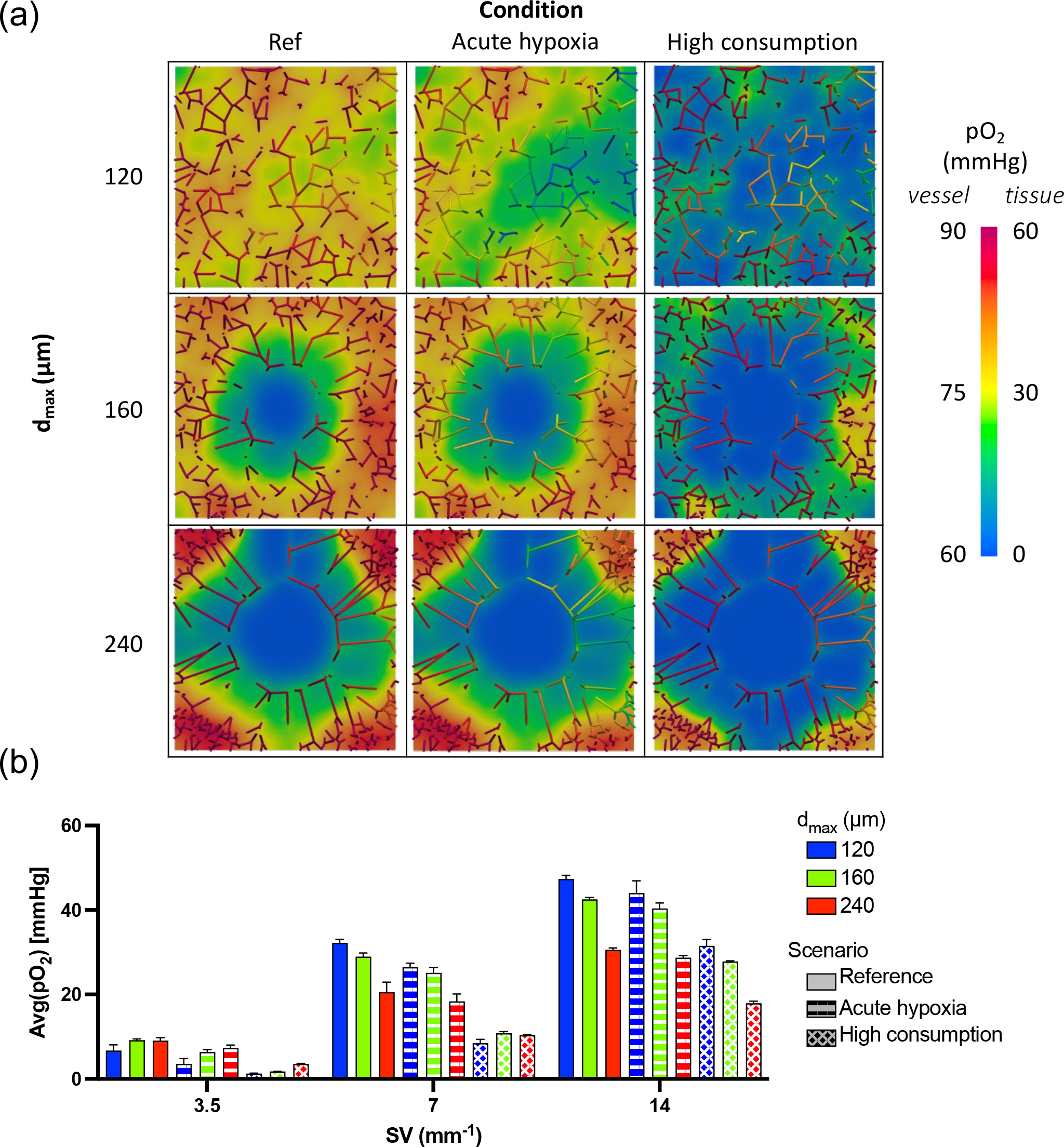
Oxygen distribution for the different oxygen scenarios. (a) Oxygen maps referring to networks with *S/V* = 7 *mm*^−1^, varying *d*_*max*_ in the vertical direction, and oxygen scenarios in the horizontal one. (b) Average oxygen partial pressure in all the 27 cases considered (3 *S/V* × 3 *d*_*max*_ × 3 *oxygen scenarios*).

Furthermore, increased oxygen consumption affects oxygenation status, creating larger hypoxic volumes with *pO*_2_ ≤ 1 *mmHg*, with a wide impact in the different microvascular conditions. To be precise, Fig. 5 shows the effect of vascular density and vascular morphology on the volume fraction of hypoxic tissue, for different vascular scenarios. We note that the hypoxic volume fraction generally decreases with vascular density and increases in perturbed vascular scenarios. Overall, hypoxia is also significantly present in the highly vascularized scenario if the morphology is not regular.

**Figure 5.**
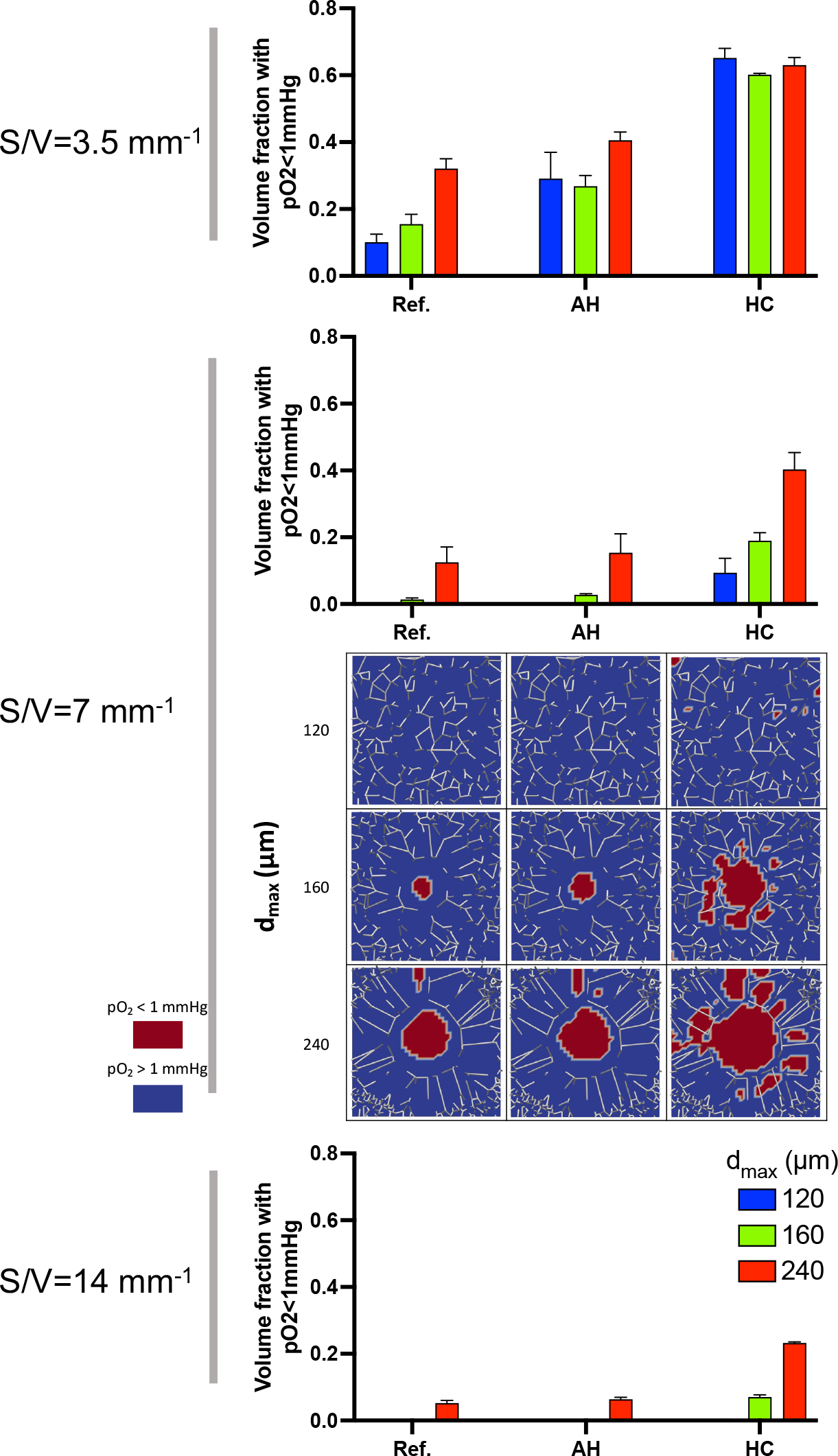
Hypoxic areas for the different oxygen scenarios. Volume fractions with *pO*_2_ ≤ 1 *mmHg* in all the 27 cases considered (3 *S/V* × 3 *d*_*max*_ × 3 *oxygen scenarios*). Plots are complemented by hypoxia maps referring to networks with *S/V* = 7 *mm*^−1^, varying *d*_*max*_ in the vertical direction, and oxygen scenarios in the horizontal one.

Furthermore, the acute hypoxia scenario is generally similar to the reference case, with substantial differences in the case of low microvascular density. Finally, the alterations observed in hypoxic volumes at low vascular density report a peculiar pattern. Regular networks are more affected than irregular networks by acute hypoxia and high consumption. Such a different behavior in low vascularized tissues agrees with data previously shown with a 2d-0d model [26]. In other words, for acute hypoxia and high consumption rates, the transition from regular to perturbed vascular morphology leaves the volume fraction of hypoxic tissue almost unchanged. In these scenarios, vascular renormalization would not improve tissue oxygenation.

### Comparing TCP in different oxygen scenarios

Using oxygen distributions, we estimate TCPs under different oxygenation scenarios (Fig. 6a-b). Taking into account photons and protons, a lower microvascular density clearly decreases the probability of successful treatment. A greatly irregular morphology (i.e., high *d*_*max*_) has a similar effect. In addition, a mid-microvasularization with mid-regularity results in better TCPs (although often unsuccessful) than irregular highly vascularized and regular low-vascularized cases. Due to its smoother response, the TIN model shows differences in the different oxygen scenarios, confirming the oxygen data (TCP under reference *>* acute hypoxia *>* high consumption). On the other hand, carbon treatment has high TCPs in all the cases considered, showing a very low dependence on microvascular characteristics.

**Figure 6.**
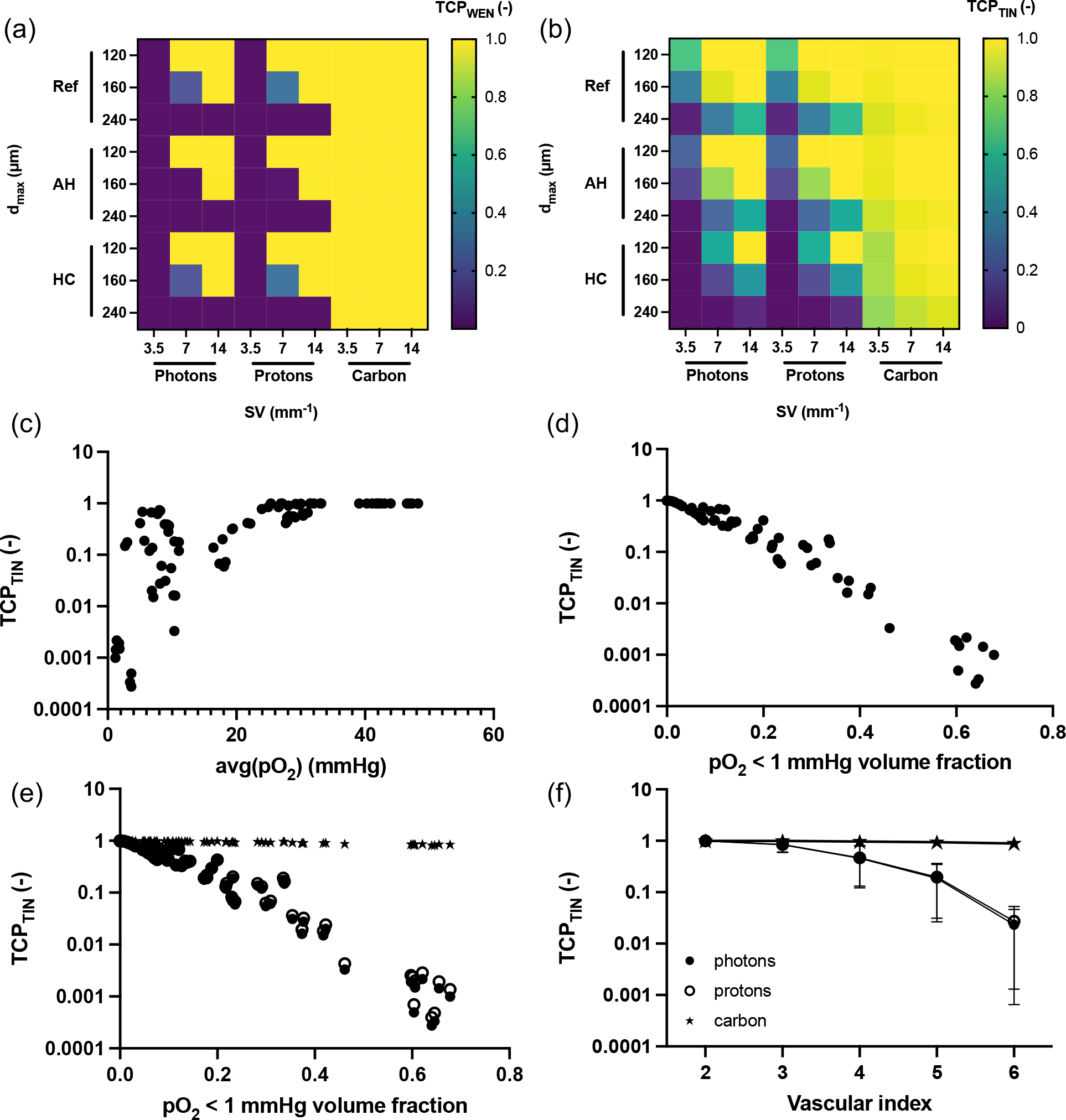
Irradiation results in all the cases analyzed. Heat maps with the TCPs for all the cases analyzed with the WEN (a) and the TIN model (b). Ref = Reference cases. AH = Acute Hypoxia cases. HC = High consumption cases. Scatterplot showing the correlation between of TCP (computed with the TIN model) with average oxygen partial pressure (c) and hypoxic volume fraction *pO*_2_ ≤ 1 *mmHg* (d). (e) Scatterplot for TCP and hypoxic volume fraction considering different treatments. (f) Effect of the *vascular index* on the TCP with different treatments.

We compare the correlations among the oxygen data and the resulting TCPs when considering photons (Fig. 6c-d). The average *pO*_2_-TCP plot shows a “safe area” depicting successful treatments (*TCP* →1) for an average oxygen partial pressure greater than 32 *mmHg*, ensuring a treatment without oxygen-related radioresistance problems. Below this threshold, the TCP is more scattered, showing how this index is not a good marker of treatment failure. On the other hand, the hypoxic volume fraction (*pO*_2_ *<* 1) is strictly correlated with the treatment outcome (*R*^2^ = .73, *p <* .0001). As also shown in the previous results, proton treatments lead to similar results when analyzing the hypoxia-TCP correlation (Fig. 6e). Carbon still shows a correlation, but with a clearly lower influence of hypoxia (coefficient of linear regression *log*_10_*TCP* - hypoxic volume fraction: photons -1.669; protons -1.673; carbon -0.247).

We note that the correlation of hypoxia with radioresistance is not surprising (see Section Radiobiological models for the radiotherapy outcome), but these results highlight the contribution to hypoxia at the microvascular scale, showing the role of microvascular morphology. To unravel the role of the microvascular network, we score the networks with two grades for the density of vascularization (1 = high, 2 = medium, 3 = low) and the morphology (1 = regular, 2 = medium, 3 = irregular). We define the vascular index as the sum of these two scores. Therefore, the lowest vascular score (= 2) represents a highly vascularized regular network. Then, the vascular score of 3 comprises a medium-vascularized regular network and a highly vascularized network with medium irregularities. With the exclusion of carbons, the TCP is close to one if the vascular index is lower than four. Indeed, at this level, the TCP starts to drop monotonically with increasing index. Interestingly, the vascular index equal to four includes heterogeneous conditions between the two indices (high-density irregular or low-density regular), which shows the importance of a comprehensive characterization of the microvascular network, both in terms of density and morphology.

## Discussion

The Biological Target Volume (BTV) includes radiobiological characteristics in the planning of radiation treatment via functional imaging. In this framework, tumor oxygenation represents a piece of essential information collected through imaging, given its importance in radiation therapy. Indeed, PET images reveal the presence of hypoxic volumes within cancer and have successfully predicted the outcome of radiation therapy [68]. However, the translation of this information into an optimized treatment plan through dose painting has not reached clinical practice [68]. The dynamic nature of hypoxia is believed to be one of the reasons for limiting dose-painting techniques, also considering that hypoxic volume changes are predictive of local recurrence [69]. A further problem lies in the so-called resolution gap. Indeed, imaging voxels are typically at the millimeter scale, whereas oxygen diffusion occurs at the micrometer scale [10]. As a consequence, very different oxygen distributions can have similar PET signals due to intra-voxel heterogeneity.

In this framework, computational models can be used to study the state of oxygenation on the microscale, as proposed in [70]. We analyzed by computational models the effect of microvascular morphology on oxygenation and, eventually, on the radiation outcome. Grogan and colleagues performed a similar study showing how the 2D and 3D representations differ when analyzing microvascular oxygen delivery and reported a negligible influence of microvascular morphology on tissue oxygenation [27]. Scott and coworkers reported a non-trivial influence of microvasculature on radiation outcome. More in detail, the morphology of the network is differently correlated with the outcome when considering high and low vascular density [26]. Their study particularly motivates the use of computational models to analyze different oxygenation scenarios. Schiavo et al. simulated an idealized tumor vasculature with heterogeneous vascular density and morphology with low vascularization toward the center of the tumor [28]. They showed that different quantities related to microvascular conditions influence tumor oxygenation and radiation outcomes in a centimeter-wide tumor with reference to clinically relevant scenarios involving both diffusion-limited and prefusion-limited hypoxia. All of these studies focus mainly on oxygen diffusion within tissue, with simplified models of oxygen transport within the microvascular network. A more complex model of oxygen transport within the microvascular network (that is, including blood flow and the presence of red blood cells [20, 52, 71, 72]) can describe the heterogeneous distribution of oxygen within vessels and its consequence on oxygen delivery, modifying the influence of microvascular morphology on radiation results.

Our study leverages a microscale oxygenation model that describes both vascular and tissue [20], applied to a domain comparable to a single imaging voxel (millimeter scale), obtaining tissue oxygen levels consistent with the literature [5, 65]. In our scenarios, we reproduced different levels of hypoxia down to a few *mmHg* (as reported by Muz and colleagues [73]), and we simulated treatments with clinically relevant schedules. We report a strong correlation between TCP and hypoxic volume (*pO*_2_ ≤ 1 *mmHg*, Figure 6). Anyway, such a correlation originates from the oxygen distribution at the microscale, reaching a subvoxel resolution. In contrast, it is lost when the resolution is limited to the size of the voxel, namely by limiting the oxygen knowledge to a single value (Figure 6). This confirms the presence of the resolution gap described by Grimes et al. [10]. Using the mean *pO*_2_ as a representative value, we can only identify a threshold required to obtain a successful treatment (*pO*_2_ |_*avg*_ ≥ 32 *mmHg*), but the outcome of treatment is highly scattered in the case of lower oxygenation.

In terms of clinical translation, these data suggest that any local de-escalation in dose painting must be approached with caution, given the current imaging capabilities. Due to resolution limitations, information available at the voxel level may hide local hypoxic spots. These areas could significantly reduce TCP, even to 10^−1^ when the average *pO*_2_ ranges from 10-20 mmHg.

Furthermore, our data show the effect of microvasculature on radiotherapy (Figures 3,6). To be more precise, a large decrease in TCP is caused independently by low vascular density or highly disordered morphology. We remark that the different vascular geometries can be representative of different tumors but also of different regions within the same tumor, as shown in [28]. Furthermore, possible combinations of vascular density, morphology, flow, and oxygen consumption lead to non-trivial results in terms of average oxygenation within the voxel (Figures 1, 4). In contrast, we do not report similar effects in hypoxic regions (*pO*_2_ ≤ 1 *mmHg*), and consequently, on TCP. These results differ from what was reported by Scott and colleagues [26]. However, a direct comparison is difficult due to the different metrics, model dimensionality, and modeled cell phenomena (e.g., cell proliferation).

In addition to classical photon-based radiation therapy, we also considered hadron therapy, which represents an alternative treatment to overcome tumor hypoxia. In fact, a higher LET is associated with a lower OER, namely, with a less pronounced oxygen-mediated radio resistance [29, 48]. In our study, we considered protons (*LET* = 4 *keV/µm*) and carbon ions (*LET* = 75 *keV/µm*). Consistently, the proton results are similar to the photon ones, whereas the carbon ions deviate toward better control of the tumor in all cases, namely, controlling hypoxic tumors. Such results agree with a similar study, in which carbon ion fractionated RT successfully controls hypoxic tumors [74]. Taking into account all of the results, we can identify cases that benefit from carbon ion treatment. In our data set, there is a clear benefit when the vascular index is greater than 4, highlighting the role of the microvasculature.

Dose enhancement represents another alternative to address hypoxia-related tumor radioresistance. The computational model proposed in this work allows us to evaluate the role of the morphology of the microvascular network in boosting the dose. We considered a further treatment, adding 10 Gy to the photon irradiation with the same fractionation schedule. Generally, boosting the dose increases TCP as expected, still less effective than carbon ion treatments (Figure 7a and b). However, the gain in TCP is often not sufficient to reach clinically acceptable values. Indeed, considering treatments with TCP ≥ 0.85, the 10 Gy boost satisfied the condition in 42% of the cases, while the photons reached 30% without the boost (Figure 7 c). Despite the increase in TCP, the boost is not effective when meeting bad microvascular morphology and, as a consequence, strong hypoxia (Figure 7d). More in detail, the gain is irrelevant with vascular index ≥ 5. Certainly, the dose can be further increased to stress this gain.

**Figure 7.**
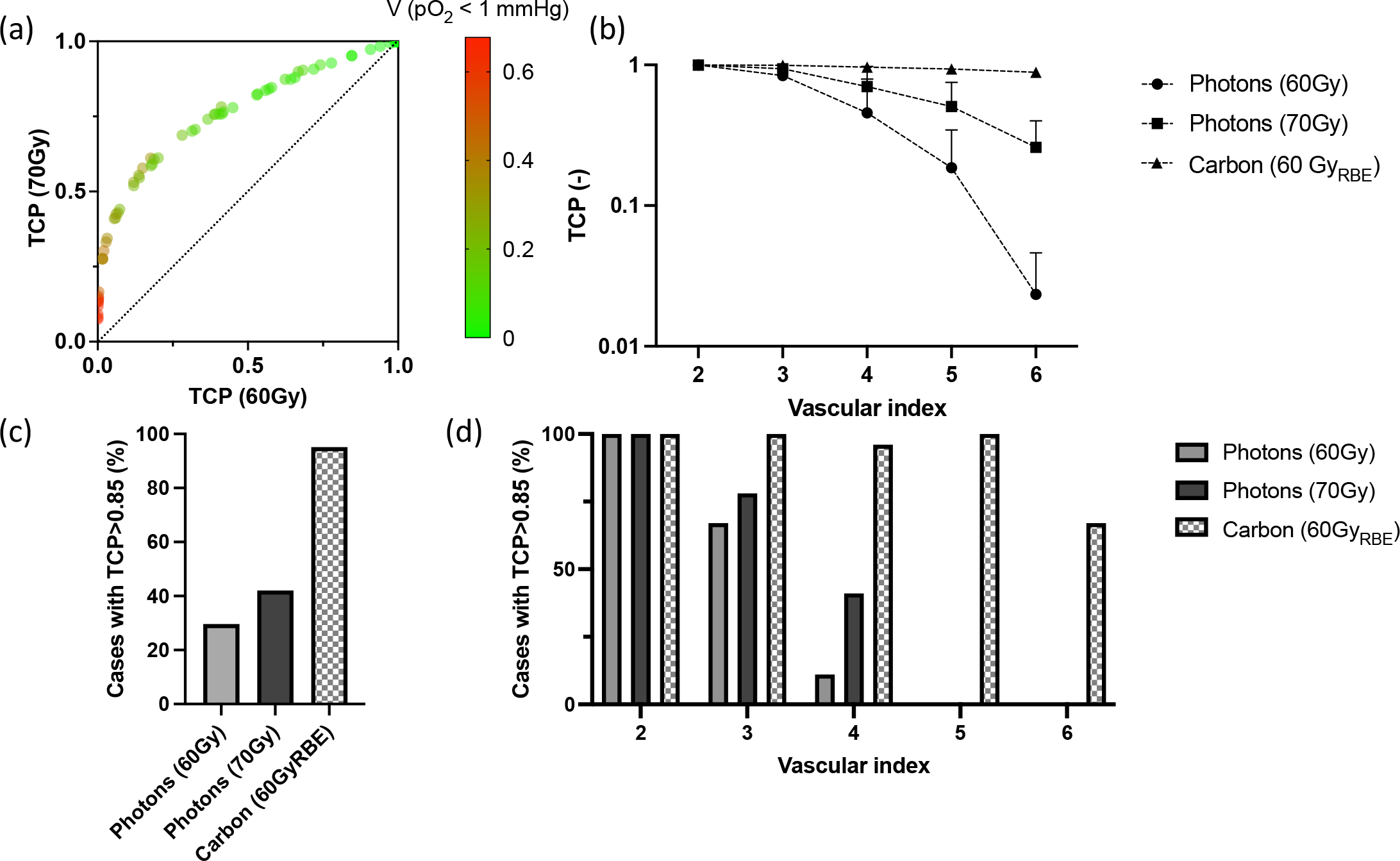
Effect of 10 Gy boost with photons. Effect of boosting with 10 Gy photons compared to 60 Gy_RBE_ treatments with photons and carbon ions. (a) Change in TCP with additional boost, colored by the hypoxic volume fraction (*pO*_2_ ≤ 1*mmHg*). Each dot represents one of the cases previously described treated with 60*Gy* or 70*Gy*. (b) TCP variations with vascular index comparing photons, boost with photons, and carbon ions. (c) Percentage of the cases with TCP ≥ 0.85 considering the three treatments applied to the entire dataset and divided by vascular index (d).

However, in clinical practice, boosting is limited by the dose administered to nearby tissues, which can induce side effects. We would like to note that, given the domain considered (namely comparable to an imaging voxel), the boosting application can also provide information on the dose painting techniques, indicating the gain achievable when the local dose increase reaches 10 Gy. In contrast, carbon ions are effective in our dataset (TCP ≥ 0.85), comprehensively in 95% of the cases. The remaining cases are mainly characterized by a vascular index equal to 6, depicting the worst vascular scenario considered in our computational data set.

We note that even if we considered a fractionated scheme, we did not include the effect of radiation damage on vascular damage (e.g., on permeability of the vascular wall and vasodilation). Vascular damage is an important factor, especially when considering high doses, but we still have limited knowledge of its mechanisms and impact on treatments [75]. Additionally, we did not consider the reoxygenation effect, namely the possible changes in oxygen consumption during treatment related to cell death. The inclusion of this effect and cell proliferation would increase the relevance of this work to the clinic. However, a detailed description of the cell number dynamics due to growth and treatment is required, both through oxygen-mediated mechanisms [76]. Furthermore, the acute hypoxia scenario can be stressed, further reducing the flow to the no-flow condition. To obtain meaningful results in such an extreme scenario, a different modeling approach must be considered involving a more complex vascular geometry, as proposed by Schiavo et al. [28]. On the other hand, we consider the average value as representative of the voxel. Different techniques can be applied and developed to provide a more physics-based extraction of the representative value (e.g., tracer binding or cellular composition [10]). An example of this application in a different field is the study by Geady et al., in which they evaluated the resolution requirements for radiomic features, considering a physics-based downsampling technique [77]. With a similar approach, we can further evaluate how a specific image (e.g. PET) is representative of the microscopic oxygen distribution.

## Conclusion

We have introduced an advanced computational approach based on mixed-dimensional modeling to investigate oxygenation in the tumor microenvironment, accounting for various conditions and vascular properties. Specifically, both vascular density and morphology play a pivotal role in oxygen distribution, thereby impacting the outcomes of radiotherapy. Using this framework, we can quantitatively assess the influence of several key factors, including the morphological characteristics of the vascular network. Interestingly, for photon and proton radiotherapy, the vascular network significantly affects treatment efficacy, while carbon-ion treatment remains effective even with irregular, low-density vessel distribution. Proton treatment does not demonstrate a distinct advantage over photon-based RT in addressing tumor hypoxia.

Our results support the existence of a resolution gap in current clinical imaging [10]. In particular, considering the potential misidentification of hypoxic regions, these findings advise caution when contemplating de-escalation in dose painting.

Furthermore, this computational tool shows promise in evaluating hypothetical scenarios and in vitro models of radiation-induced damage to the microvasculature [78].

## Supporting information

### Appendix. S1 Parameter estimation for radiobiological models

This supplementary section describes the procedure to fit parameters for the Tinganelli model (TIN) based on data published by Wenzl et al. The TIN model reads as follows:

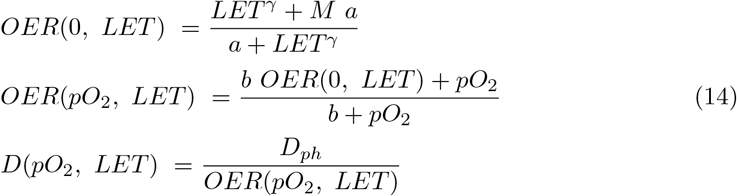

where *M, a, γ*, and *b* are parameters fitted to experimental data.

We first estimate the parameters *M, a*, and *γ* by RMSE minimization against the data reported in Figure 1 (solid line) by Wenzl et al. [48] (Fig. S3). This data set represents the OER-LET relation under no oxygen conditions (*p*_*O*_2 = 0.01*mmHg* ≃0.001%). Therefore, we use this data set to estimate *OER*(0, *LET*). The parameter values are *M* = 2.81, *a* = 522.45 *keV/µm*, and *γ* = 1.38.

As a second step, using the data in Figure 3 of the same work, we estimated the parameter *b* (Fig. S3). We run the RMSE minimization on the data for *LET* = 2 *keV/µm*. Consequently, the fit is less accurate for *LET* = 100 *keV/µm*. The resulting value for *b* is 1.24 *mmHg*.

Finally, we report the *OER* function considering photon, protons, and carbon, namely *OER*_*photons*_(*pO*_2_, 2 *keV/µm*), *OER*_*protons*_(*pO*_2_, 4 *keV/µm*), and *OER*_*carbon*_(*pO*_2_, 75 *keV/µm*) (Fig. S3).

### Appendix. S2 Numerical discretization and solvers

The discretization of the problem ℱ + ℋ + 𝒪 is achieved using the finite element method. One of the main advantages of our formulation is that the computational meshes of Ω and Λ are completely independent. For this reason, we address the two approximations separately.

We denote with 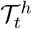 an admissible family of partitions of 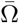 into tetrahedrons *K*

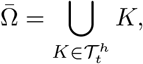

that satisfies the usual conditions of a conforming triangulation of Ω. Here, *h* denotes the characteristic size of the mesh, that is, 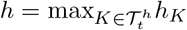, where *h*_*K*_ is the diameter of the simplex *K*. Moreover, we implicitly assume that Ω is a *polygonal* domain. The solution of ℱ is approximated using discontinuous piecewise-polynomial finite elements for pressure and ***H***_*div*_-conforming *Raviart-Thomas* finite elements for velocity, namely

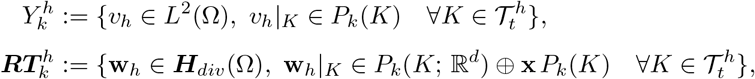

for every integer *k* ≥ 0, where 𝒫_*k*_ indicates the standard space of polynomials of degree ≤ *k* in the variables **x** = (*x*_1_, …, *x*_*d*_). The lowest order *Raviart-Thomas* approximation has been adopted, corresponding to *k* = 0 above.

Concerning the capillary network, we adopt the same approach used at the continuous level, that is, we split the network branches into separate subdomains. Furthermore, each curved branch Λ_*i*_ is approximated by a piecewise linear 1D line, denoted with 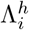. More precisely, the latter is a partition of the i-th network branch made up of a sufficiently large number of segments, named 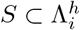. In this way, we obtain the following discrete domain:

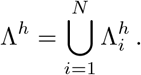

The solution of ℱ on a given branch 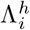 is approximated by using continuous piecewise polynomial finite element spaces for both pressure and velocity. Since we want the vessel velocity to be discontinuous at multiple junctions, we define the related finite element space over the whole network as the collection of the local spaces of the single branches. Conversely, the pressure has been assumed to be continuous over the network; therefore, its finite element approximation is standard. We will use the following families of finite element spaces for pressure and velocity, respectively:

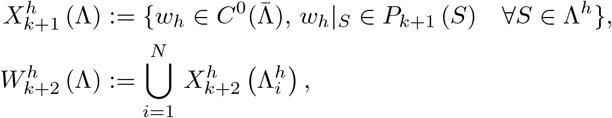

for every integer *k* ≥ 0. As a result, we use generalized Taylor-Hood elements on each network branch, satisfying in this way the local stability of the mixed finite element pair for the network. At the same time, we guarantee that the pressure approximation is continuous across the entire network Λ^*h*^. In particular, for the numerical experiments shown later, we have used the lowest order, that is, *k* = 0.

For hematocrit we proceed as for the velocity approximation. In particular, we approximate equation H with the finite element space 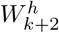 defined on 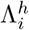. For the sake of generality, let us define the families of discrete subspaces of the functional spaces for *k* ≥ 0:

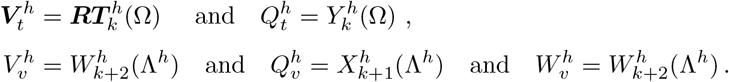

According to the work presented in [18, 20], to which we remand for the meaning of all the symbols of the equations that follow, the discrete equations for the microvascular flow and red blood cell transport (the ℱ+ ℋ model) are the following: find 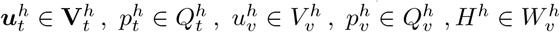 such that

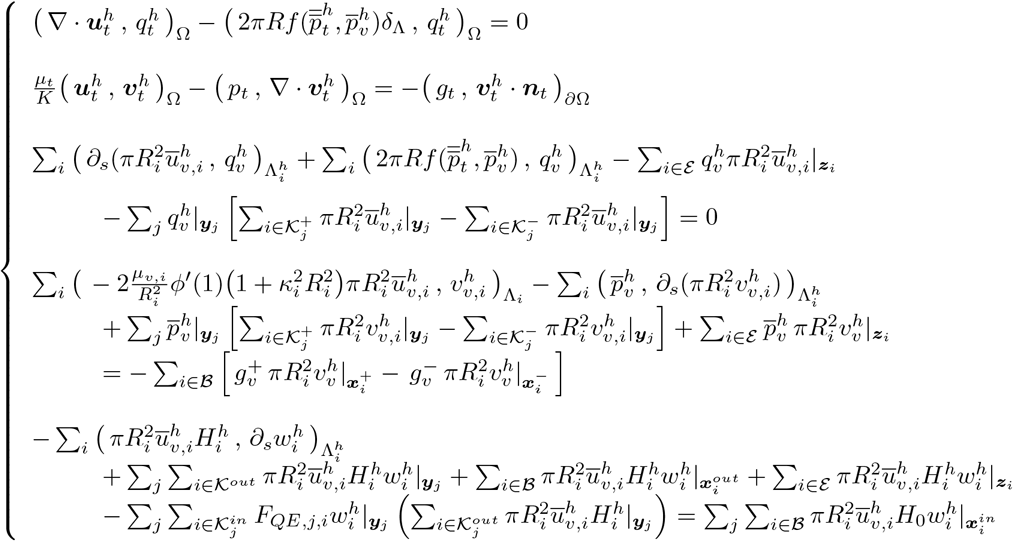

Proceeding similarly, the discrete formulation of the oxygen transport problem, namely problem 𝒪, is obtained projecting the weak formulation of the equations in suitable discrete finite element spaces.

Let us define 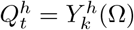 and 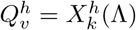 for k ≥ 0 where 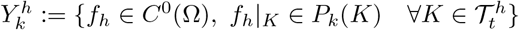, for every integer *k* ≥ 0, where *P*_*k*_ indicates the standard space of polynomials of degree ≤ *k* in variables **x** = (*x*_1_, …, *x*_*d*_) and 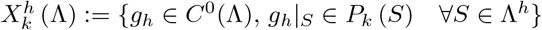, for every integer *k* ≥ 0.

We also apply a fixed-point method to linearize the system of equations by evaluating the reaction term in the tissue and the oxyhemoglobin concentration in the microvessels at the previous iteration. More precisely, we define the new coefficient, Ψ^(*k*−1)^ as the oxyhemoglobin concentration at the previous iteration:

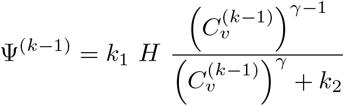

Such formulation of the oxyhemoglobin term highlights the effect of the transport of free oxygen with respect to the bound-hemoglobin oxygen. Then, the blood velocity can be re-written as follows:

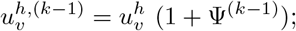

Then, the discrete equations for the oxygen transport model become: find 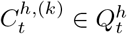 and 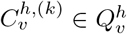 such that,

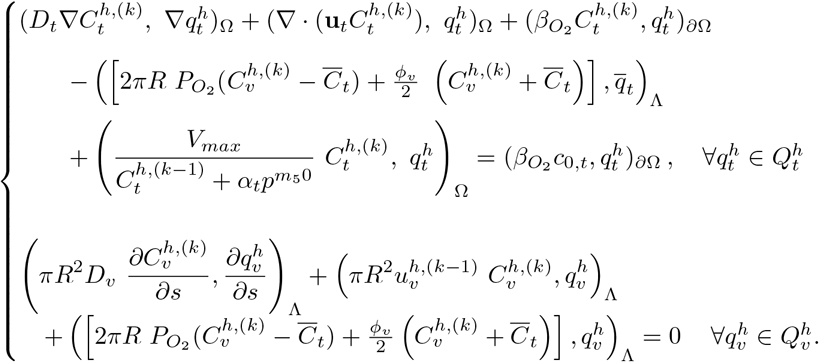

As mentioned above, the meaning of all variables and symbols of these equations is described in [79].

In order to calculate the numerical solution of the problem, let us introduce the algebraic formulation of the complete problem. The linearization of the problem is beneficial; in fact, at each stage of the fix-point technique a linear system is generated and must be solved. To be more concise, we report the full formulation of the discrete system only for the oxygen transport problem 𝒪, remanding to the interested reader the discrete version of ℱ+ ℋ to [18].

For problem 𝒪 the number of degrees of freedom (DOF) of the discrete spaces 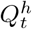 and 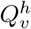 is 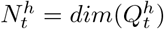 and 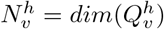 and the finite element basis functions are 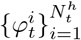 for 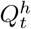 and 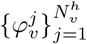 for 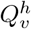. Then the numerical solution for the oxygen concentration can be written as a linear combination of these base functions 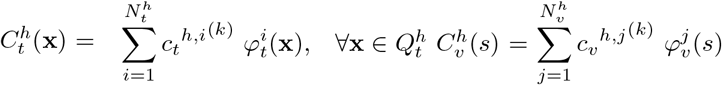. Substituting these expressions in the weak discrete problem and exploiting the linearity of the inner product, we obtain the following linear system for each iterative step:

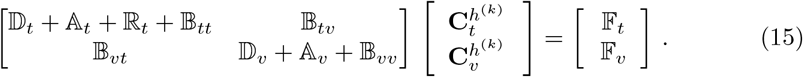

The submatrices and subvectors in (15) are defined as follows:

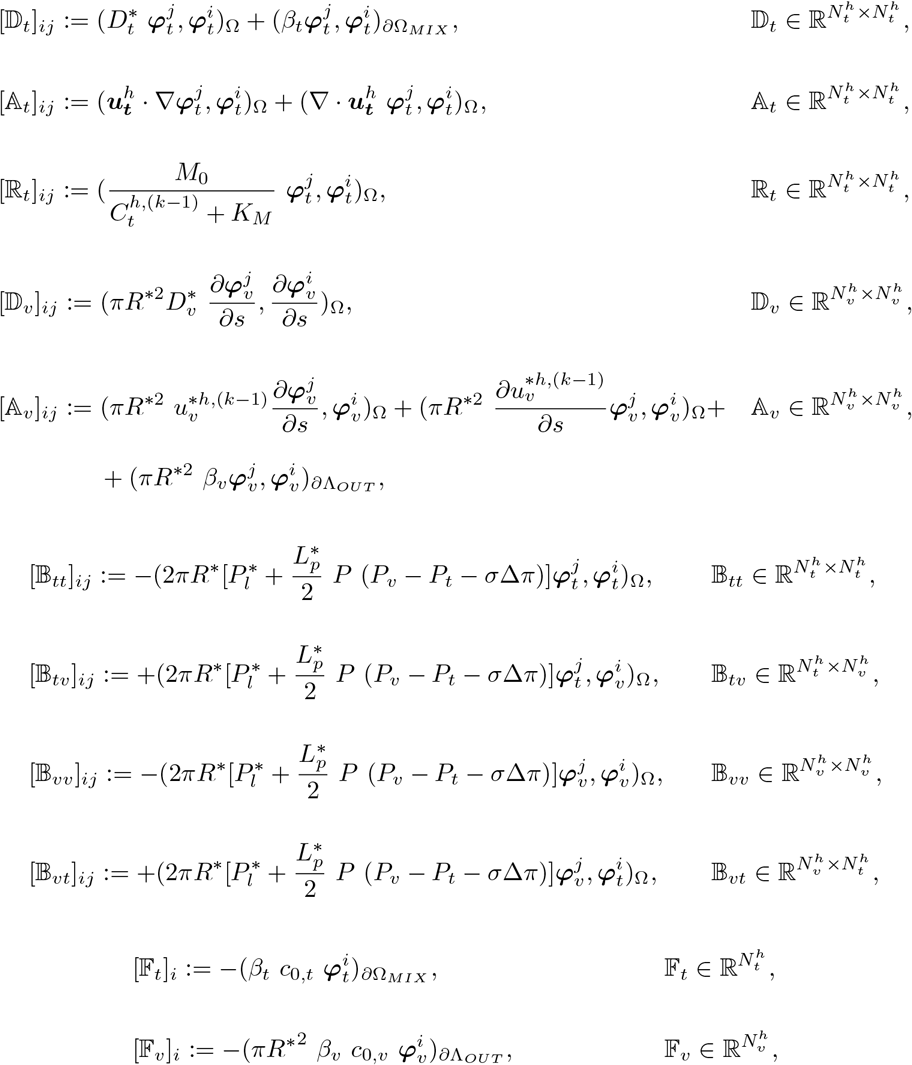

Where 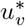 is:

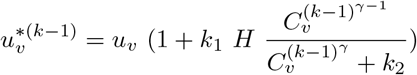

## Supplementary Images

**Supplementary Figure S1.**
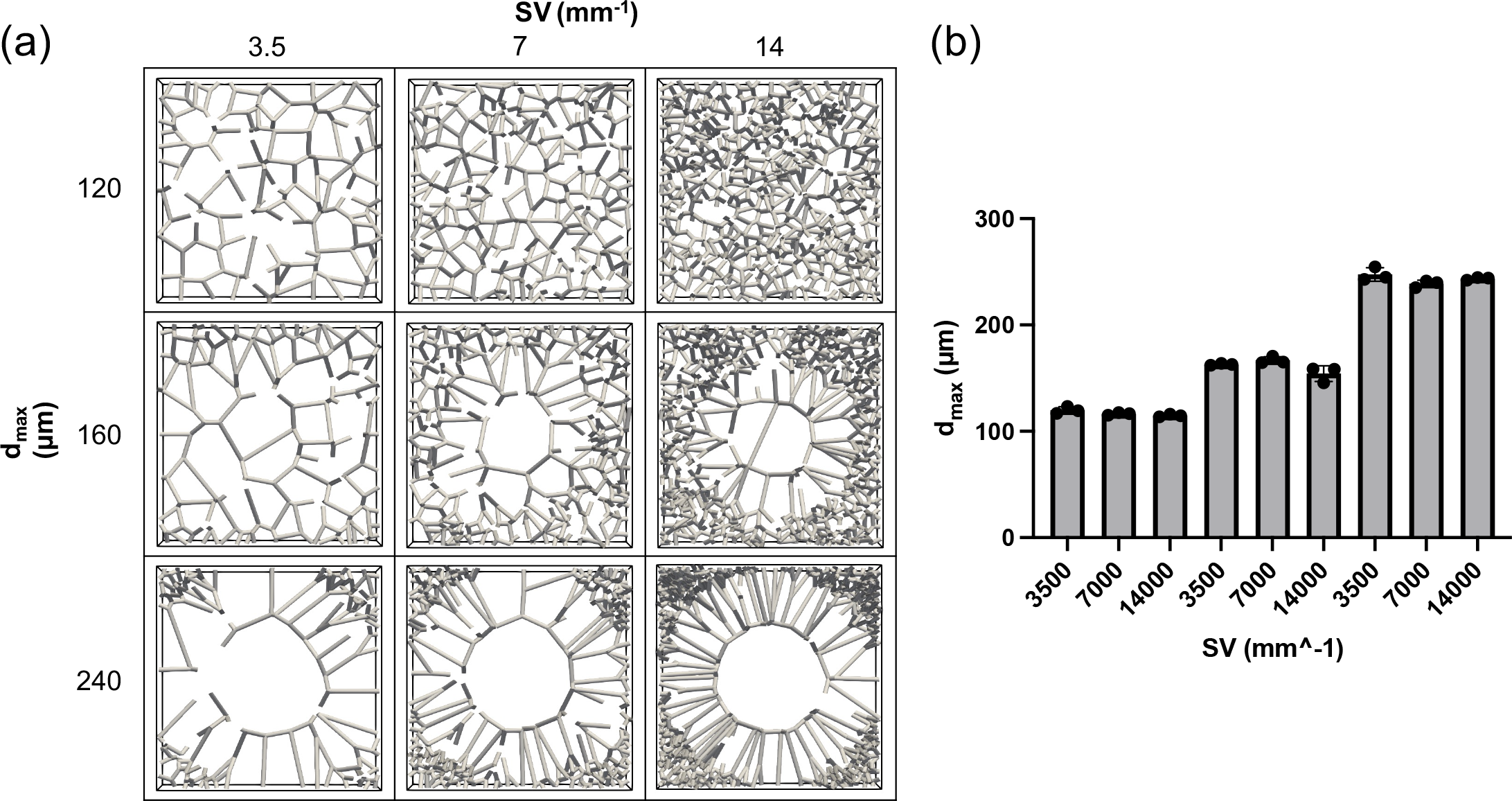
Set of networks with data. Representative images of the networks generated on the tissue slab. (a) One replicate per condition, showing the effect of the two metrics *S/V* and *d*_*max*_. (b) Evaluation of *S/V* and *d*_*max*_ for each networks.

**Supplementary Figure S2.**
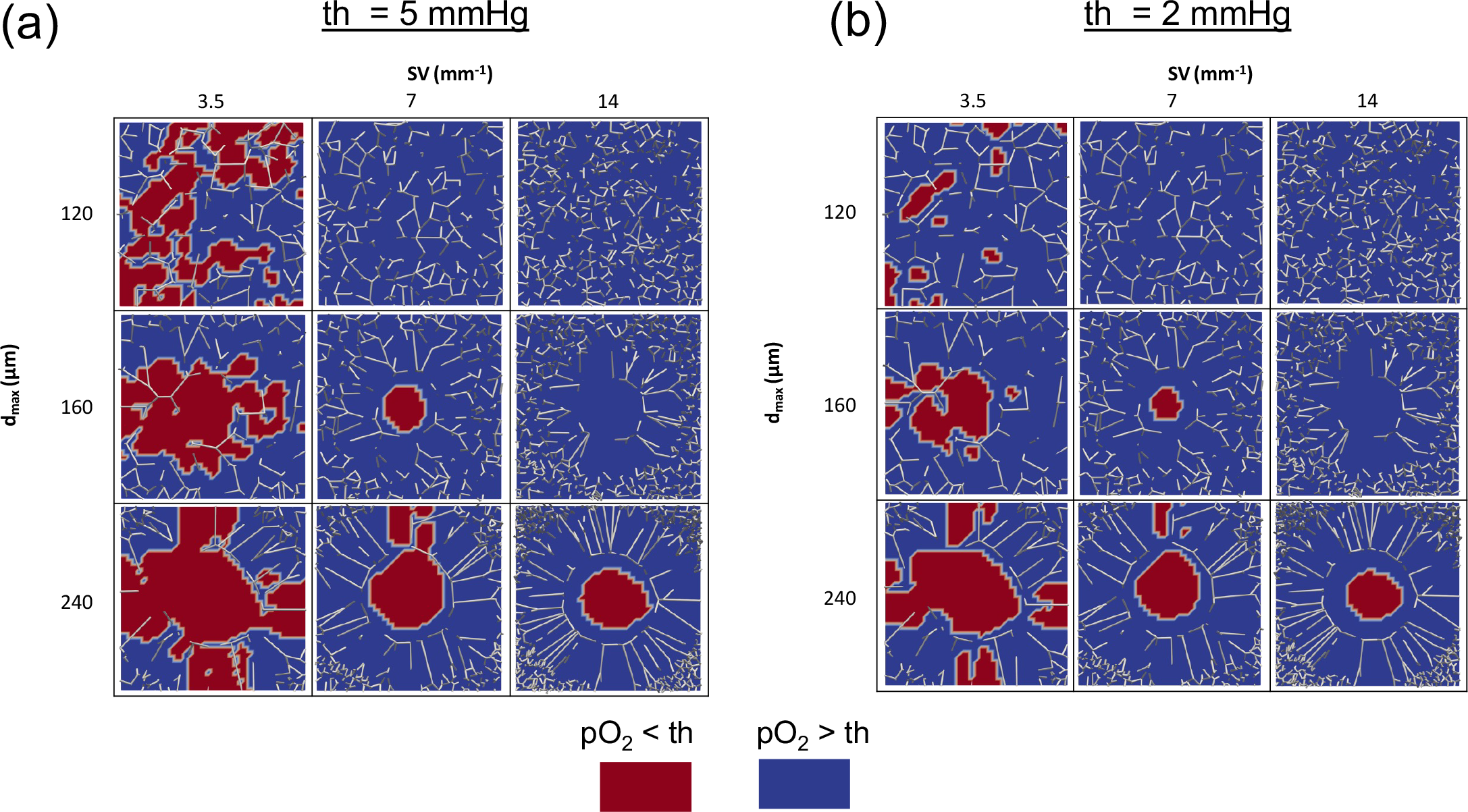
Spatial distribution of low oxygen areas in the reference scenario. Low oxygen areas in the nine cases with (a) *th* = 5*mmHg* and (b) *th* = 2*mmHg*.

**Supplementary Figure S3.**
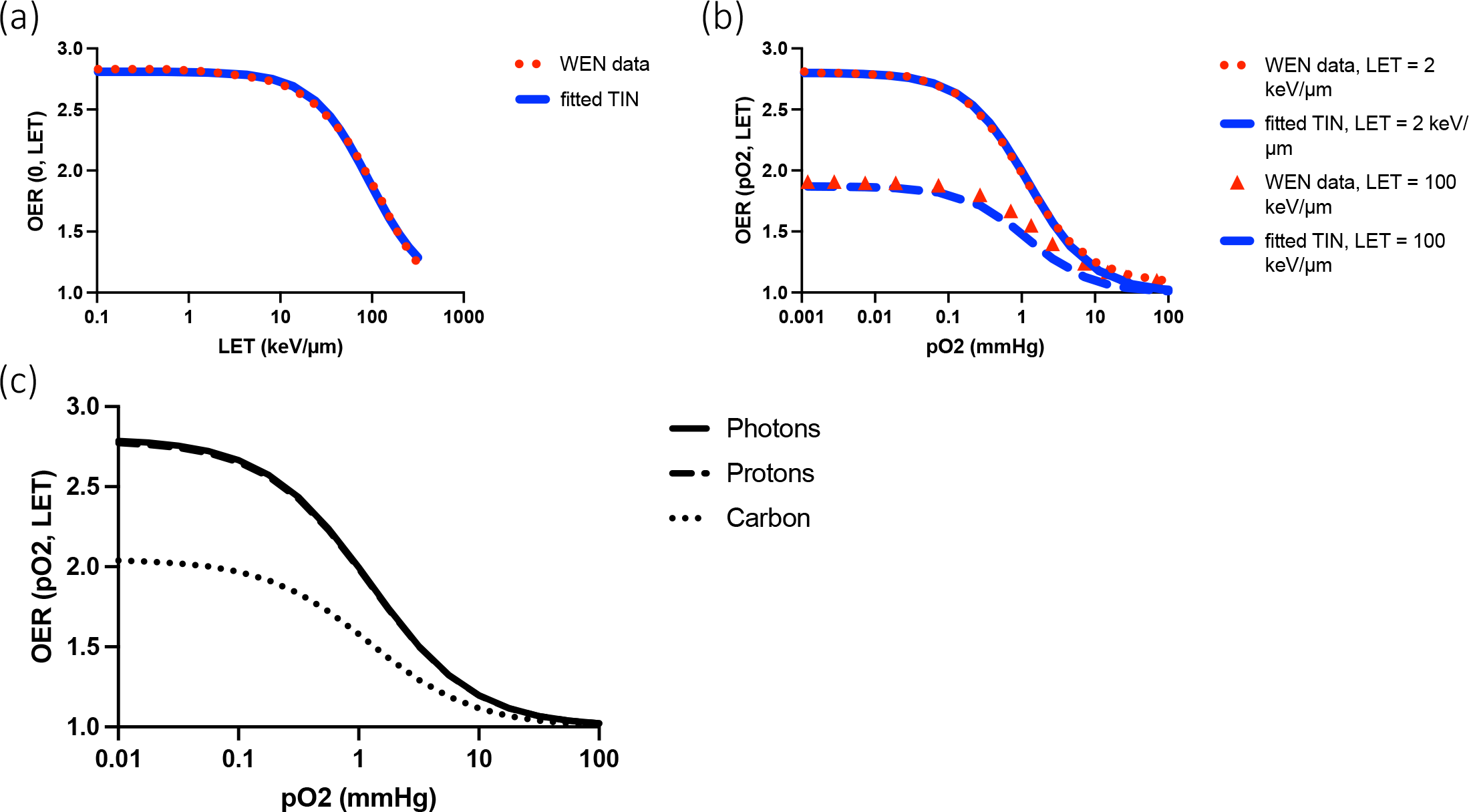
TIN model fit against WEN data. TIN model fitting against WEN data. (a) Fitting for the *OER*(0, *LET*) function. (b) Fitting for the *OER*(*pO*_2_, *LET*) relation with *LET* = 2 *keV/µm* and *LET* = 100 *keV/µm*. (c) The resulting OER model for photons, protons, and carbon ions.

## Acknowledgments

This work was supported by the AIRC Investigator Grant, no. IG21479 (TZ is the PI of the study). The funders had no role in study design, data collection and analysis, decision to publish, or preparation of the manuscript.

